# Decreased spliceosome fidelity inhibits mTOR signalling and promotes longevity via an intron retention event

**DOI:** 10.1101/2022.03.07.483153

**Authors:** Wenming Huang, Chun Kew, Stephanie A. Fernandes, Anna Loerhke, Lynn Han, Constantinos Demetriades, Adam Antebi

## Abstract

Changes in splicing fidelity are associated with loss of homeostasis and ageing^1–3^, yet only a handful of splicing factors have been shown to be causally required to promote longevity ^1–3^, and the underlying mechanisms and downstream targets in these paradigms remain elusive. Surprisingly, we found a hypomorphic mutation within RNP-6/PUF60, a spliceosome component promoting weak 3’ splice site recognition, which causes aberrant splicing, elevated stress responses, and enhances longevity in *Caenorhabditis elegans*. Through genetic suppressor screens, we identify a gain-of-function mutation within *rbm-39*, an RNP-6 interacting splicing factor, which increases nuclear speckle formation, alleviates splicing defects and curtails longevity caused by *rnp-6* mutation. By leveraging the splicing changes induced by RNP-6/RBM-39 activities, we uncover a single intron retention event in *egl-8*/phospholipase C B4 as a key splicing target prolonging life. Genetic and biochemical evidence show that neuronal RNP-6/EGL-8 downregulate mTORC1 signaling to control organismal life span. In mammalian cells, PUF60 downregulation also potently and specifically inhibits mTORC1 signaling. Altogether, our results reveal that splicing fidelity modulates mTOR signaling and suggest a potential therapeutic strategy to delay ageing.

## Main

PUF60 (poly-U-binding factor 60 kDa) encodes an essential splicing factor that binds uridine (U)-rich tracts and promotes association of the U2 small nuclear ribonucleoprotein complex (U2 snRNP) with primary transcripts ^4,5^. PUF60 is required for cell viability, proliferation and migration *in vitro*. Its deficiency in patients causes developmental defects ^6–8^ and overexpression is associated with tumorigenesis ^9,10^, but a role in metabolism and ageing is completely unknown. In a previous genetic screen for *C. elegans* longevity regulators using cold tolerance as proxy, we had identified a novel mutation in the worm ortholog of PUF60, *rnp-6*, carrying a Gly281Asp substitution (referred to as *rnp-6(G281D)* hereafter) in the second RNA recognition motif (RRM), which alters resistance to multiple abiotic stresses and extends life span ^11^.

We decided to characterize the nature of this mutation in more detail and found that *rnp-6(G281D)* behaved as a recessive, hypomorphic allele, since (1) *rnp-6(G281D)*/+ heterozygotes were as cold sensitive as *wild-type* controls (Extended Data Fig. 1a); (2) Knock-down of *rnp-6* by RNAi bacterial feeding (*rnp-6i*) enhanced cold tolerance in the *wild-type* background and caused developmental arrest in *rnp-6(G281D)* mutants (Extended Data Fig. 1b); (3) Overexpression of *rnp-6(wt)* but not *rnp-6(G281D)* transgene fully reversed *rnp-6(G281D)* cold tolerance phenotype (Fig. 1a). Moreover, *rnp-6(wt)* transgene also fully abolished the longevity phenotype (Fig. 1b). To characterize the cellular function of *rnp-6*, we tagged endogenous *rnp-6* with GFP using CRISPR/Cas9. Consistent with its role as an essential splicing factor, *rnp-6(wt)* was ubiquitously expressed in all examined tissues and mainly localized in the nucleus (Extended Data Fig. 1c). GFP tagged endogenous *rnp-6(G281D)* showed a similar expression pattern, but was present at significantly lower levels (Fig. 1c and Extended Data Fig. 1d), which was also validated by Western blot (Extended Data Fig. 1e). RNAseq analysis also showed that *rnp-6(G281D)* caused changes in mRNA processing and transcription similar to but not as extensive as *rnp-6i*, including alternative splicing, intron retention, circular RNA formation (Fig. 1d–e, Extended Data Fig. 1f–i, Supplementary Table 1-3), as well as differential gene expression (Extended Data Fig. 1j, Supplementary Table 4), confirming that *rnp-6(G281D)* represents a reduction-of-function mutation. Strikingly, ~80% of the differentially expressed genes (DEG) (1142 out of 1366 genes in *rnp-6(G281D)* and 3730 out of 4707 genes in *rnp-6i*) were upregulated (Supplementary Table 4). Gene ontology analysis of the differentially expressed genes showed that stress response was among the most enriched physiological categories in both *rnp-6(G281D) and rnp-6i* (Extended Data Fig. 1k, Supplementary Table 5), suggesting that impaired spliceosome function triggers cellular stress responses.

**Figure 1.**
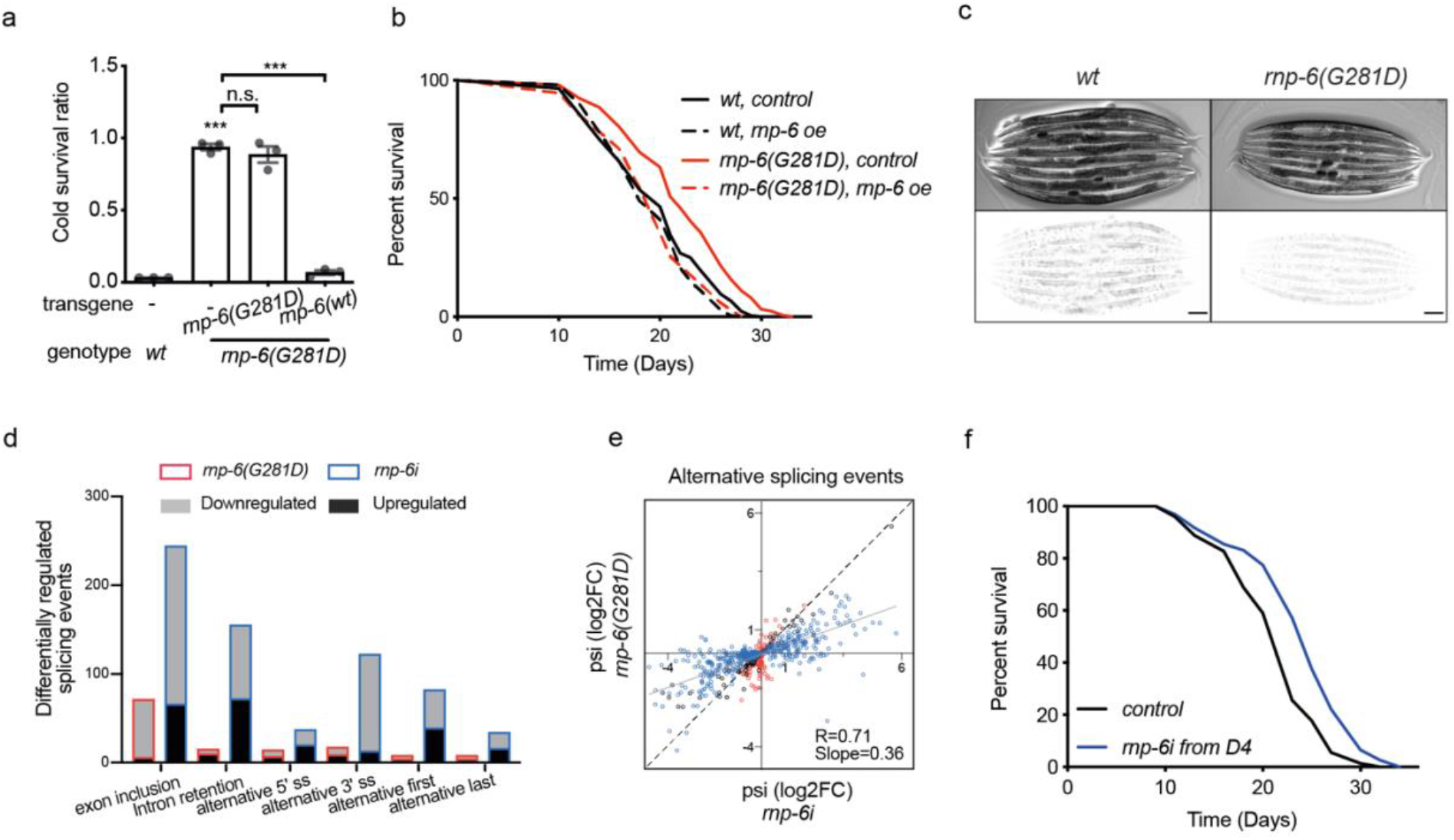
Reduction of *rnp-6* function promotes longevity. a, Effect of transgenic *rnp-6* over-expression on rescue of cold tolerance. n=3. Mean ± SEM. ***p < 0.001. One-way ANOVA. b, Effect of transgenic *rnp-6* over-expression on rescue of life span. n=2. For all life span experiments, survival curves depict one representative experiment. Other repeats are shown in Supplementary Table 16. c, Expression of endogenous *rnp-6(wt) and rnp-6(G281D)* tagged with GFP imaged in young adult stage worms. Scale bar, 100 μm. Top and bottom panels represent DIC and GFP fluorescent images, respectively. Fluorescence is inverted to show better contrast. d, Quantity of differentially regulated alternative splicing changes found in *rnp-6(G281D)* and *rnp-6i*. e, Correlation of *rnp-6(G281D)* and *rnp-6i*-induced alternative splicing changes. Each dot represents the log2-transformed fold changes of an event relative to *wild-type* control. Blue, red and grey dots indicate the events which are significantly changed by *rnp-6i*, *rnp-6(G281D)* and both, respectively. f, Survival assay of *rnp-6i* treatment from day 4 adulthood. n=4.

Next, we asked if *rnp-6i* mimicked *rnp-6(G281D)* longevity. To bypass developmental defects, we performed *rnp-6* RNAi knockdown during adulthood. Whereas *rnp-6i* in day 1 adults (coinciding with the onset of reproduction) decreased life span (Extended Data Fig. 1l), *rnp-6i* initiated from day 4 adult (coinciding near the end of reproduction) onward significantly extended it (Fig. 1f). These results show that, like several other essential genes ^12,13^, knockdown can be detrimental early in life but beneficial later, and imply that the fine-tuning of *rnp-6* activity is critical for longevity.

In order to dissect the functional network underlying *rnp-6* longevity, we performed unbiased genetic suppressor screens. Notably, we observed that *rnp-6(G281D)* exhibited a temperature sensitive (*ts*) growth phenotype, which could be fully rescued by *rnp-6(wt)* overexpression (Fig. 2a, Extended Data Fig. 2a, b). We reasoned that crucial regulators which suppress this *ts* defect could also alleviate other *rnp-6(G281D)* functions. We screened ~20,000 genomes and isolated 13 mutants (Extended Data Fig. 2c). Using Hawaiian SNP variant mapping, whole genome sequencing and CRISPR/Cas9 gene editing, we succeeded in identifying two candidates: *rnp-6(dh1187)* and *rbm-39(dh1183)* (Fig. 2b). The *rnp-6(dh1187)* intragenic mutation led to a glutamate to lysine substitution (*rnp-6(E161K)*), which corresponds to E188 in human PUF60 (Extended Data Fig. 4a, b). This residue mediates interdomain RRM1-RRM2 contacts in the PUF60 crystal structure ^14^, and may affect salt bridge formation. RBM-39 encodes an RNA binding protein, whose human ortholog, RBM39, functions as a splicing factor and is involved in early spliceosome assembly ^15^. Similar to PUF60, RBM39 contains two central RRM domains and a C-terminal U2AF-homology motif (UHM) domain, but additionally harbors an N-terminal arginine–serine-rich (RS) domain (Fig. 2c) implicated in nuclear speckle formation ^16^. The *rbm-39(dh1183)* mutation caused a serine to leucine substitution (S294L) in the second RRM (Fig. 2c). This residue is conserved in nematodes, but changed to proline in higher organisms (Extended Data Fig. 2d, e). Interestingly, a proline to serine substitution at this same position in human RBM39 changes its conformation and renders resistance to the anti-cancer drug, indisulam, an aryl sulfonamide that facilitates RBM39 proteasomal degradation ^17^, highlighting the pivotal role of this residue in regulating RBM39 function.

**Figure 2.**
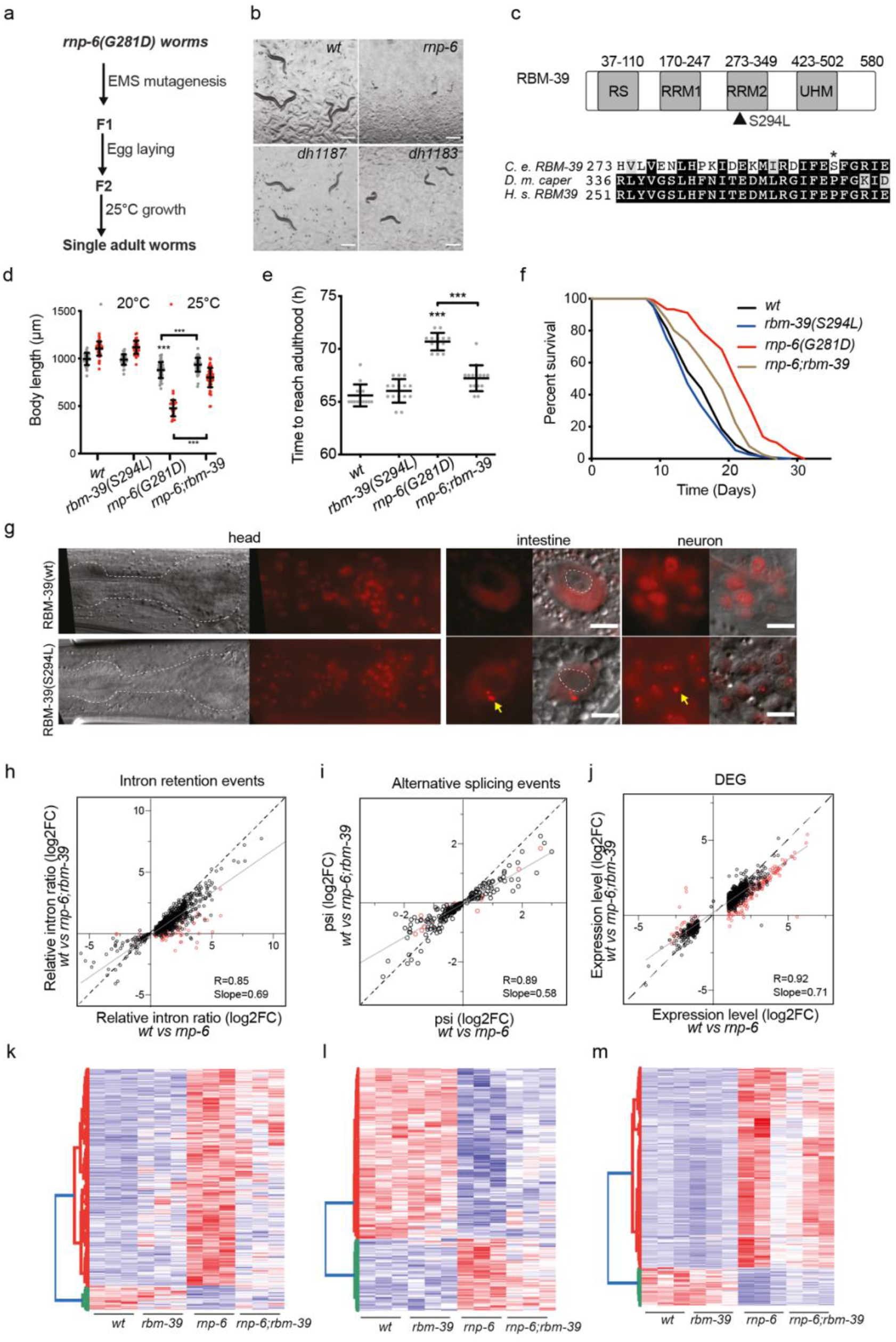
*rbm-39* functionally interacts with *rnp-6*. a, Schematic workflow of the suppressor screen. b, Representative images of *dh1183* and *dh1187* suppressors grown at the restrictive temperature 25°C. Scale bar, 500 μm. c, Protein domain structure of RBM-39 and sequence alignment of RBM-39 homologs. Filled triangle and asterisk denote the location of the Serine 294 to Leucine mutation. d-f, Effect of *rbm-39(S294L)* on *rnp-6(G281D) ts* (d) developmental rate (e) and longevity (f) phenotypes. n=3. d and e, Mean ± SD. ***p < 0.001. One-way ANOVA. g, Nuclear localization of RBM-39. Yellow arrows indicate RBM-39(S294L) intranuclear puncta. Scale bars, 5 μm. h-m, Effect of *rbm-39(S294L)* on intron retention (h and k), alternative splicing (i and l) and gene expression changes (j and m) as shown by scatter plots and heat maps. Each dot represents the log2-transformed fold changes of an event relative to *wild-type* control. Red dots denote the events which are significantly suppressed by *rbm-39(S294L)*.

*rmb-39(S294L)* represents a semi-dominant allele as *rmb-39(S294L)/+* heterozygotes partially suppressed the *rnp-6(G281D) ts* phenotype (Extended Data Fig. 2f). To further clarify *rmb-39* function, we tested the effect of decreased *rbm-39* activity on *rnp-6(G281D) ts* phenotype. *rbm-39* RNAi (*rmb-39i*) exacerbated *rnp-6(G281D)* phenotypes, and caused a further decrease in body size, yet had little effect on *wild-type* controls (Extended Data Fig. 2g, h). Similarly, another reduction-of-function allele *rbm-39(R251C) ^18^* further delayed developmental rate and decreased body size at 20°C, and caused complete embryonic lethality at 25°C (Extended Data Fig. 2i). These results suggest that *rbm-39* and *rnp-6* function in concert, and support the notion that the *rbm-39(S294L)* suppressor likely defines a specific change- or gain-of-function allele.

Since *rbm-39(S294L)* largely reversed the *ts* defect (Fig. 2d), we next asked if it also suppressed other *rnp-6(G281D)* phenotypes visible at the permissive temperature (20°C). Notably, *rbm-39(S294L*) significantly restored body size (Fig. 2d), developmental rate (Fig. 2e), infection tolerance (Extended Data Fig. 2j) and decreased *rnp-6(G281D)* life span (Fig. 2f), but had little effect on its own (Fig. 2d–f), suggesting that *rbm-39(S294L*) ameliorates *rnp-6(G281D)* function.

In order to address the potential mechanisms behind *rbm-39(S294L))*-mediated suppression of the *rnp-6(G281D)* phenotypes, we first examined if *rbm-39(S294L)* restored reduced *rnp-6(G281D)* protein levels. Western blot experiments, however, showed that *rbm-39(S294L)* had no impact on either *rbm-39* or *rnp-6* protein levels (Extended Data Fig. 3a, b). We then wondered if *rbm-39(S294L)* altered the subcellular localization of RNP-6 or RBM-39. Endogenously mKate2-tagged RBM-39(WT) and RBM-39(S294L) were ubiquitously expressed and found mainly within the nucleus of various cell types, similar to RNP-6 (Fig. 2g, Extended Data Fig. 3c). Interestingly, we observed that the RBM-39(S294L) mutant protein, but not RBM-39(WT), formed prominent discrete puncta within the nucleus without altering RNP-6 localization (Fig. 2g, Extended Data Fig. 3d). These puncta resembled nuclear speckles implicated in regulating transcription and splicing ^19,20^. Time lapse imaging showed that these puncta were, like other nuclear speckles, highly dynamic (Extended Data Video 1). In addition, RBM-39 and RNP-6 mutually co-immunoprecipitated (Extended Data Fig. 3e), suggesting they associate in a complex. These results imply that *rbm-39(S294L*) might alleviate *rnp-6(G281D)* defects through enhanced splicing activity. To test this hypothesis, we performed RNA sequencing analysis with *wt*, *rnp-6(G281D), rbm-39(S294L)* and *rnp-6;rbm-39* double mutants. In accord with our idea, we observed that *rbm-39(S294L)* altered the transcriptional profile (Extended Data Fig. 3f) and decreased total circular RNA and intron reads of *rnp-6(G281D*) mutant (Extended Data Fig. 3g-h, Supplementary Table 6-7). Further analysis revealed that *rbm-39(S294L)* significantly suppressed a subset of the intron retention (115/954 events) (Fig. 2h, k, Supplementary Table 7), alternative splicing (19/251 events) (Fig. 2i, l, Supplementary Table 8), and differential gene expression (275/1709 events) changes (Fig. 2j, m, Supplementary Table 9) caused by *rnp-6(G281D*), and globally trended towards alleviating many such events. Gene ontology enrichment analysis showed that the *rnp-6* dependent DEGs suppressed by *rbm-39(S294L)* were significantly enriched in the stress response category (Extended Data Fig. 3i), indicating that this process might be associated with longevity. These results confirm that *rbm-39(S294L)* ameliorates *rnp-6(G281D)* splicing activity.

In addition, we found that the *rnp-6(E161K)* intragenic mutation was also a potent suppressor of *rnp-6(G281D)*, which fully restored all measured phenotypes to *wild-type* levels (Extended Data Fig. 4c-e). It also significantly suppressed mRNA processing as well as transcriptional changes (Extended Data Fig. 4f-k, Supplementary Table 10-13), confirming that, like *rmb-39* mutation, restoration of splicing correlates with reversal of phenotype.

To decipher the downstream mechanisms by which RNP-6/RBM-39 complex regulates longevity, we focused on splicing events. In particular, intron retention is an important but not well-understood mechanism of gene expression regulation ^21^. It is most associated with down-regulation of gene expression via nonsense mediated decay ^22^ and has recently emerged as an important splicing feature in both normal aging and longevity interventions ^1,23,24^. To reveal functionally relevant targets for the RNP-6/RBM-39 complex, we focused on intron retention induced by *rnp-6(G281D)*, and restored by *rbm-39(S294L)*. We narrowed down the list of candidates to 44 events by cross-referencing with the *rnp-6(E161K)* revertant and manual curation in the genome browser (Supplementary Table 14). These 44 events correspond to 42 genes and notably all showed increased intron retention in *rnp-6(G281D)*. We performed RNAi knockdown to screen for their impact on *wild-type* life span, reasoning that both RNAi and intron retention should result in partial loss-of-function. Of those genes tested, we found one candidate, *egl-8*, whose knockdown yielded significant life span extension in *wild-type*, but did not further extend *rnp-6(G281D)* longevity (Fig. 3a). We further confirmed this genetic interaction with an *egl-8(n488)* null allele (Fig. 3b). *egl-8* encodes an ortholog of human phospholipase C beta 4 (PLCB4). It plays vital physiological roles in neurotransmission ^25,26^, life span and infection response in *C. elegans* ^27,28^, though the underlying molecular mechanisms are not well understood. RNAseq data indicated that *rnp-6(G281D)* specifically increased the retention of intron 8 of *egl-8* (Extended Data Fig. 5a, b), while showing only a minor effect on total mRNA expression level (Extended Data Fig. 5c). RT-PCR results validated that the intron retention phenotype was significantly suppressed by *rbm-39(S294L)* (Fig. 3c, d) and *rnp-6(E161K)* (Extended Data Fig. 5d, e). Intron 8 harbors a weak non-canonical splice acceptor site (Extended Data Fig. 5f), consistent with a role of PUF60 in weak 3’ splice site recognition ^4^, and its retention introduces a premature stop codon in the transcript (Extended Data Fig. 5g), which could either result in mRNA degradation by nonsense-mediated mRNA decay or give rise to a non-functional truncated protein. To examine expression, we tagged endogenous EGL-8 with mNeonGreen at the N-terminus. mNeonGreen tagged EGL-8 was mainly detected in head neurons as well as intestinal adherens junctions (Extended Data Fig. 5h), in agreement with previous immunofluorescence staining results ^25^. As expected, the expression levels of EGL-8 in neurons and the nerve ring were significantly lower in *rnp-6(G281D)* compared with *wild-type* controls (Fig. 3e, f). Furthermore, neuronal expression of *rnp-6(+)* or the fully spliced *egl-8(+)* cDNA largely suppressed *rnp-6(G281D)* longevity (Fig. 3g, h). These findings are consistent with the idea that *rnp-6(G281D)* promotes longevity via intron retention of *egl-8* within the nervous system.

**Figure 3.**
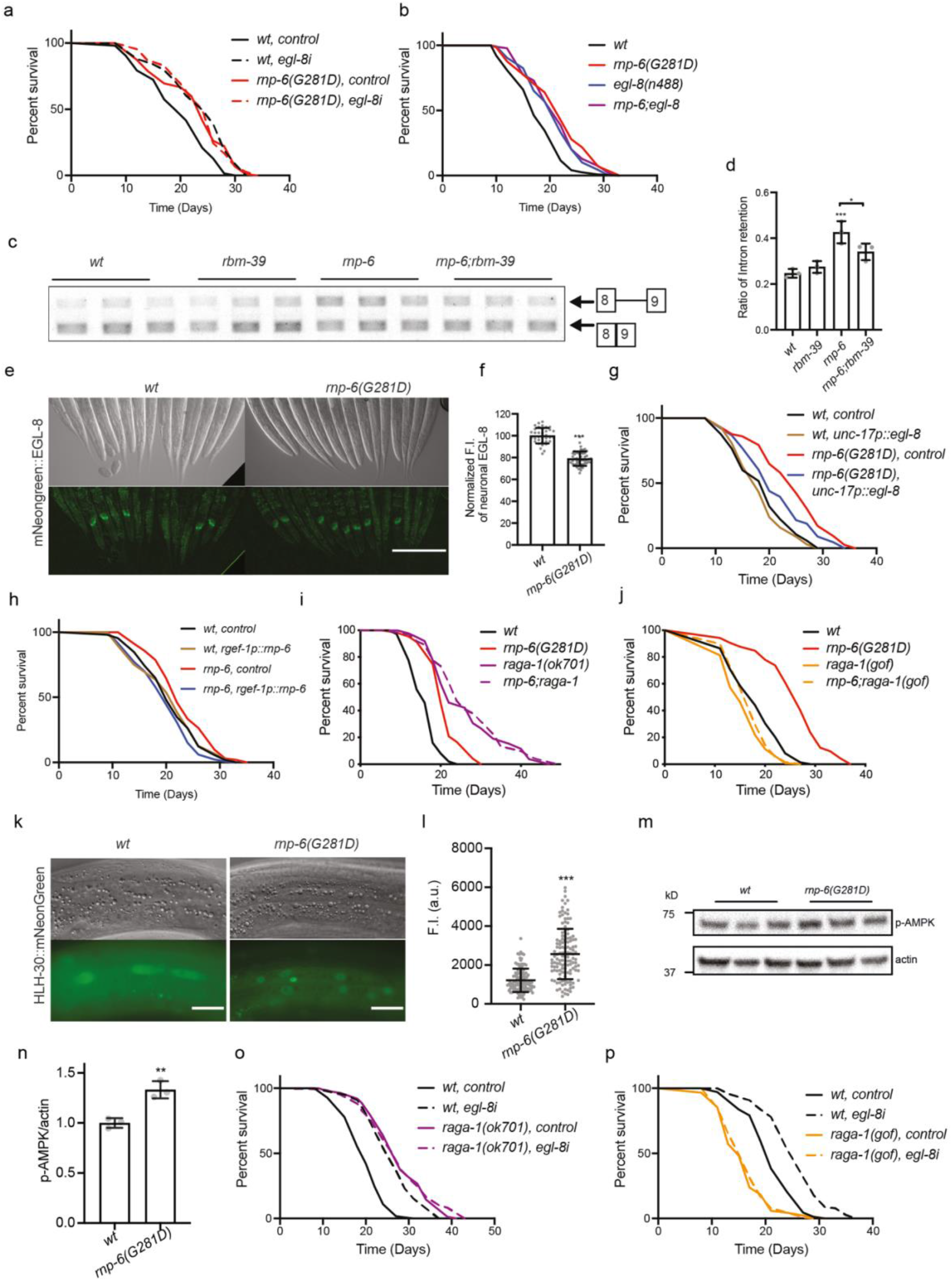
*rnp-6(G281D)* inhibits mTORC1 signaling via *egl-8* intron 8 retention. a, Life span of *egl-8* RNAi knock down on *rnp-6(G281D)* mutant. n=3. b, Life span of *egl-8* null mutation on *rnp-6(G281D)* mutant. n=3. c-d, Effect of *rbm-39(S294L)* on *egl-8* intron 8 retention. Mean ± SEM. *p < 0.05, ***p < 0.001. One-way ANOVA. e-f, Effect of *rnp-6(G281D)* on EGL-8 expression. Scale bars, 200 μm. g, Life span analysis of *egl-8* cDNA neuronal expression. n=3. h, Life span analysis of *rnp-6* cDNA neuronal expression. n=3. i-j, Effect of *raga-1* loss-of-function *(ok701)* and gain-of-function *(gof)* on *rnp-6(G281D)* life span. n=3. k-l, *rnp-6(G281D)* alters HLH-30/TFEB nuclear localization. n=3. Mean ± SD. ***p < 0.001. unpaired t-test. m-n, *rnp-6(G281D)* alters AAK-2/AMPK phosphorylation. n=3. Mean ± SEM. **p < 0.01. unpaired t-test. o-p, Effect of *egl-8 RNAi* on *raga-1* loss-of-function and gain-of-function mutants life span. n=3

To identify potential signaling pathways in which *rnp-6* might act, we performed genetic epistasis analysis, first focusing on two major conserved longevity pathways: reduced insulin/IGF (*daf-2*, insulin/IGF receptor) and mTORC1 inhibition (*raga-1*, core component of the lysosomal amino acid sensing machinery ^29^). We found that *raga-1* but not *daf-2* mutant mediated longevity was non-additive with *rnp-6* (Fig. 3i, Extended Data Fig. 6a), suggesting that *rnp-6*(*G281D)* might work in the same pathway as mTORC1. In accord with this view, the *raga-1* gain-of-function transgene, which shortens *wild-type* worm life span ^30^, completely abolished *rnp-6(G281D)* longevity (Fig. 3j), suggesting that *rnp-6* functions upstream of *raga-1* to promote mTORC1 signaling activity. Loss-of-function mutations in transcription factors FOXO/DAF-16 and HSF1/HSF-1, which mediate the output of reduced mTORC1 longevity ^31,32^, also completely abrogated *rnp-6(G281D)* longevity (Extended Data Fig. 6b, c). Furthermore, *rnp-6(G281D)* longevity was also non-additive with dietary restriction (Extended Data Fig. 6d), another longevity regime inhibiting mTORC1 ^33^. To obtain further evidence, we tested whether molecular outputs of mTORC1 signaling were also altered in the *rnp-6(G281D)* mutant. Downregulation of mTORC1 signaling results in enhanced nuclear accumulation of HLH-30/TFEB (transcription factor EB) ^34,35^ and increased levels of phosphorylated AAK-2/AMPK (AMP-activated protein kinase) ^36^. Consistently, we observed significant increase in both of HLH-30 nuclear localization (Fig. 3k, l) and AAK-2 phosphorylation (Fig. 3m, n) in *rnp-6(G281D)* mutants. Altogether, our results indicate that *rnp-6(G281D)* inhibits mTORC1 signaling activity through *raga-1*.

Since EGL-8 serves as a downstream target of *rnp-6*, we wondered if it also interacts with mTORC1 signaling pathway in regulating longevity. Interestingly, *egl-8(n488)* loss-of-function mutation significantly inhibited mTORC1 activity as indicated by increased HLH-30 nuclear localization (Extended Data Fig. 6e, f) and AMPK phosphorylation (Extended Data Fig. 6g, h). Furthermore, *egl-8i* did not further extend the life span of *raga-1 null* mutants (Fig. 3o), while *raga-1* gain-of-function transgene completely suppressed *egl-8i-*induced longevity (Fig. 3p). These results demonstrate that *egl-8* acts upstream of *raga-1*, linking *rnp-6* to mTORC1 signaling, and are consistent with previous studies showing that phospholipases can control mTORC1 activity via the generation of phosphatidic acid ^37^.

Last, we sought to understand if the functional interaction of RNP-6 and mTORC1/RAGA-1 was evolutionarily conserved. To this end, we knocked down PUF60 by siRNA in HEK293FT cells, and measured various outputs of mTORC1 signaling. Because *rnp-6* regulates mTORC1 upstream of *raga-1* (Fig. 3i, j), we presumed that PUF60 may be affecting amino acid signaling to mTORC1. In accord with our hypothesis, PUF60 knockdown decreased mTORC1 reactivation upon amino acid re-supplementation, assayed by the phosphorylation of its direct substrates S6K (ribosomal protein S6 kinase beta-1) and TFEB (Fig. 4a), without influencing mTORC2 activity (assayed by Akt phosphorylation) (Extended Data Fig. 6i). Accordingly, we consistently observed decreased Raptor (regulatory-associated protein of mTOR) protein levels upon PUF60 knockdown, while the levels of the respective mTORC2 core component, Rictor, or of mTOR itself, were largely unaffected (Fig. 4a, Extended Data Fig. 6i). In line with the *C. elegans* results, and further supporting decreased mTORC1 activity, PUF60 knockdown enhanced the nuclear localization of TFE3 transcription factor (Fig. 4b, c). Because amino acid sufficiency controls mTORC1 localization to lysosomes via promoting Raptor binding to the lysosomal Rag GTPase dimers, we then hypothesized that PUF60 may be regulating mTORC1 activity by influencing its subcellular localization. Indeed, knocking down PUF60 caused a significant drop in the colocalization of mTOR with the lysosomal marker LAMP2 (lysosome-associated membrane glycoprotein 2) in cells re-supplemented with amino acids (Fig. 4d, e). These findings reveal that PUF60 acts as a specific and integral part of the mTORC1 signaling pathway, influencing the amino-acid-induced activation of mTORC1 at the lysosomal surface.

**Figure 4.**
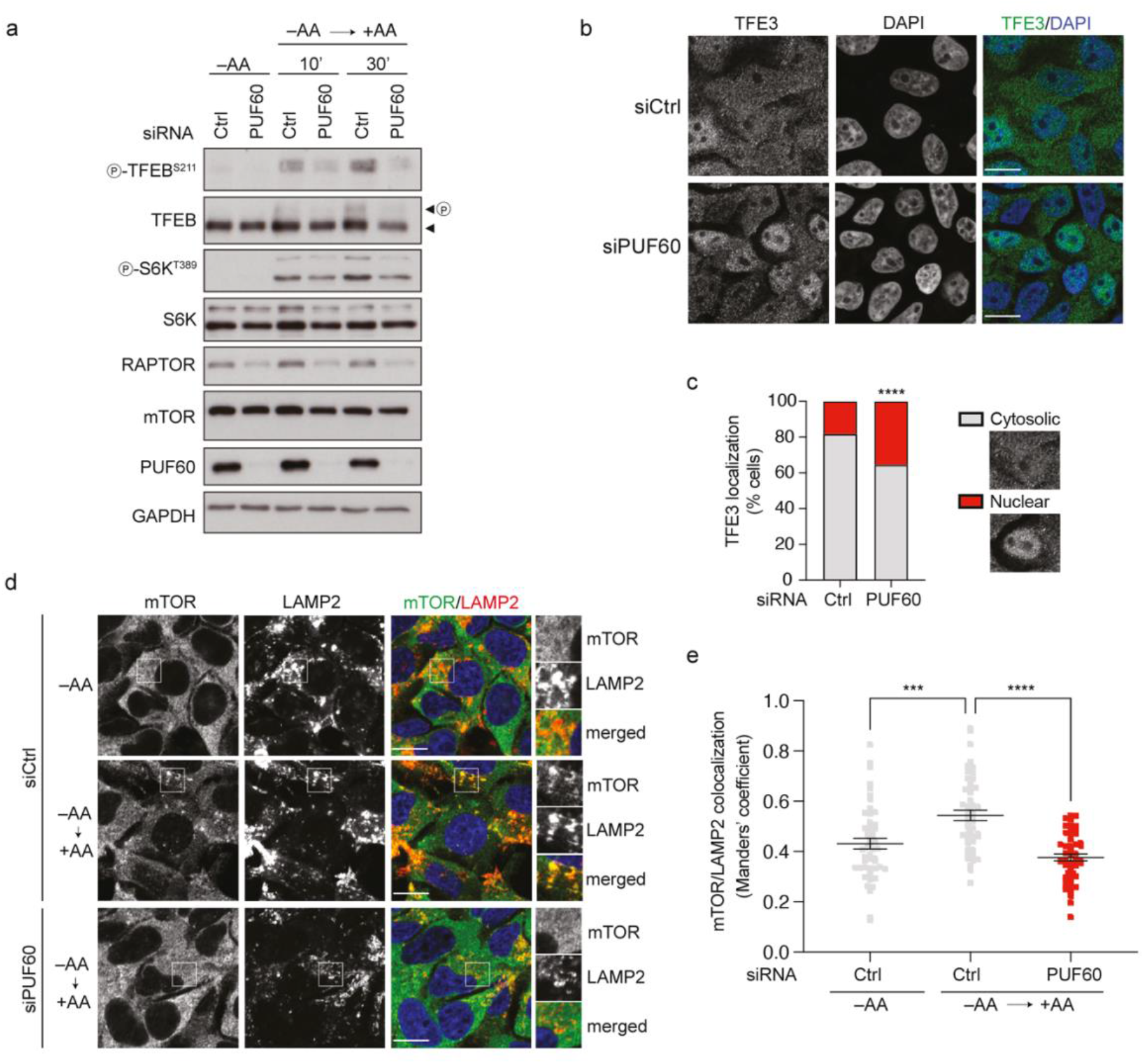
PUF60 regulates mTORC1 signaling in mammalian cells. a, Effect of PUF60 knock down on mTORC1 activity. Arrowheads indicate bands corresponding to different protein forms, when multiple bands are present. P: phosphorylated form. n = 3. b-c, PUF60 knock down alters nuclear TFE3 localization. Scale bars, 10 μm. n = 3. ****p<0.0001, two-way ANOVA. d-e, PUF60 knock down alters lysosomal localization of mTORC1. Magnified insets shown to the right. Scale bars, 10 μm. n = 3. mean ± SEM. ns, not significant, ***p < 0.001, ****p<0.0001, one-way ANOVA.

Our evidence indicate that RNP-6 and RBM-39 intimately work together to impact splicing fidelity to regulate mTOR signaling and longevity (Extended Data Fig. 6j). Though the detailed biochemical mechanisms remain to be elucidated, it is intriguing that both splicing factors contain a similar domain architecture and are associated with U2AF complexes involved in 3’ splice site selection ^16^, pinpointing this step as critical to fidelity regulation *in vivo*. Conceivably such events serve as sensors of endogenous or environmental stress linked to spliceosome activity, intron retention and RNA processing ^38^. Importantly, their action upstream of mTOR signaling may provide novel approaches to manipulate this pathway in ageing, metabolism and disease. Precise targeting of PUF60, and perhaps RBM39, could be used to downregulate mTORC1 signaling to confer health benefits similar to rapamycin and other rapalogs ^39,40^. Conversely, as many spliceosomeopathies that reduce spliceosomal function trigger growth defects ^6,41–44^, it may be possible to treat these diseases with mTOR modulators.

## Methods

### *C. elegans* strains and maintenance

The following strains were used in this study: N2 (*wild-type*), *rnp-6(dh1127)*, *rbm-39(syb1074)*, *rnp-6(dh1127);rbm-39(syb1074)*, *rbm-39(gk454899)*, *rnp-6(dh1127);rbm-39(gk454899), (wt, dhEx1139 dhEx1139[rnp-6p∷FLAG∷HA∷GFP∷rnp-6b cDNA∷unc-54 3′UTR, myo-3∷mcherry])*, *(rnp-6(dh1127), [rnp-6p∷FLAG∷HA∷GFP∷rnp-6b cDNA∷unc-54 3′UTR, myo-3∷mcherry]), (wt, dhEx1147[rnp-6p∷FLAG∷HA∷GFP∷rnp-6b(G281D) cDNA∷unc-54 3′UTR*, *myo-3∷mCherry], (rnp-6(dh1127), dhEx1147[rnp-6p∷FLAG∷HA∷GFP∷rnp-6b(G281D) cDNA∷unc-54 3′UTR, myo-3∷mCherry], (wt, dhEx1208[unc-17p∷gfp∷egl-8, myo-3p∷mCherry]), (rnp-6(dh1127), dhEx1208[unc-17p∷gfp∷egl-8, myo-3p∷mCherry])*, *egl-8(syb3661), rnp-6(dh1127);egl-8(syb3661), raga-1(ok701)*, *rnp-6(dh1127);raga-1(ok701), egIs12[raga-1(gf); Pofm-1∷GFP], (rnp-6(dh1127), egIs12[raga-1(gf); Pofm-1∷GFP]), daf-2(e1370), rnp-6(dh1127);daf-2(e1370), daf-16(mu86), rnp-6(dh1127);daf-16(mu86)*. All mutant strains obtained from CGC or NBRP were outcrossed with our N2 at least twice before experiments. Worms were maintained at 20°C following standard procedures ^45^. For all experiments, synchronization of the animals was done by the egg laying.

### Cell culture and treatments

All cell lines were grown at 37°C, 5% CO_2_. Human female embryonic kidney HEK293FT (#R70007, Invitrogen; RRID: CVCL_6911) cells were cultured in high-glucose DMEM (#41965039, Thermo Fisher Scientific), containing 10% Fetal Bovine Serum (FBS) and 1% Pen/Strep. The cells were purchased from Invitrogen before the initiation of the project. Their identity was validated by the Multiplex human Cell Line Authentication test (Multiplexion GmbH), which uses a single nucleotide polymorphism (SNP) typing approach, and was performed as described at www.multiplexion.de. All cell lines were regularly tested for *Mycoplasma* contamination, using a PCR-based approach and were confirmed to be *Mycoplasma*-free.

### Plasmid construction and transgenesis

For *rnp-6* rescue plasmid*, rnp-6* promoter (3135 bp) was amplified from N2 genome and inserted into pDC4 vector generate *rnp-6p∷FLAG∷HA∷GFP∷∷unc-54 3′UTR* construct. Then, *rnp-6b* cDNA was amplified from N2 cDNA and cloned into this plasmid to generate *rnp-6p∷FLAG∷HA∷GFP∷rnp-6b cDNA∷unc-54 3′UTR* rescue plasmid. Site-directed mutagenesis (Q5® Site-Directed Mutagenesis Kit, NEB) was performed to incorporate G281D point mutation to generate *rnp-6p∷FLAG∷HA∷GFP∷rnp-6b(G281D) cDNA∷unc-54 3′UTR* plasmid. In order to generate neuronal rescue plasmid, *rnp-6* promoter was replaced by neuronal-specific promoter *rgef-1 (2670 bp). unc-17p∷gfp∷egl-8* plasmid is a kind gift from Stephen Nurrish (Harvard Medical School). The microinjection experiments were performed according to standard protocol ^46^. 10 ng/μl plasmid of interest together with 5 ng/μl co-injection marker plasmid (*myo-3p∷mCherry*) were injected to gonad of young adult stage worms. Positive offspring were singled to maintain stable lines. PCR primers related to these plasmids are available in Supplementary Table 15.

### EMS mutagenesis screen and mapping

The suppressor screen was done with *rnp-6(G281D)*. L4 larvae were exposed to 0.15% ethyl methane sulfonate (EMS, Sigma) in M9 buffer for 4h at room temperature. After recovery overnight, young adult P0 adult animals were transferred to new plates for egg laying at 20°C. After 3 days growing, adult F1 worms were bleached and eggs were seeded on NGM plates and incubated at 25°C. After 3 days, single adult F2 worms from the plates and maintained the mutant worms at 20°C. Hawaiian-SNP mapping and whole genome sequence was used to map the causative mutation ^47^. *rnp-6(G281D)* mutation was firstly introduced to Hawaiian CB4856 by outcrossing 6 times. Then, the EMS mutants were crossed with Hawaiian males which carry *rnp-6(G281D)* mutation. Eggs of F1 generation worms were growing at 25°C and adult F2 were singled after three days. The heat resistant strains were then pooled together, and genomic DNA were purified using Gentra Puregene Kit (Qiagen). The pooled DNA was sequenced on an Illumina HiSeq platform (paired-end 150 nucleotide). MiModD pipeline (http://www.celegans.de/en/mimodd) was used to identify the mutations. The WS220/ce10 *C. elegans* assembly was used as reference genome for annotation. The causative mutations were either confirmed by CRISPR/Cas9 or multiple outcross.

### Protein alignments

T-Coffee algorithm ^48^ was used to align RNP-6, RBM-39 and their homologs from different species. Protein sequences of *H. sapiens* RBM39 (Uniprot: Q14498), *C. elegans* RBM-39 (Uniprot: Q9N368) and *D. melanogaster* Caper (Uniprot: Q9VM49) were used in Figure 2D. Protein sequences of *H. sapiens* PUF60 (Uniprot: Q9UHX1), *C. elegans* RNP-6 (Uniprot: Q9N3S4) and *D. melanogaster* Hfp (Uniprot: Q8T6B9) were used in Figure S2B. Protein sequences of *C. brenneri* RBM-39 (Uniprot: G0NLU2), *C. elegans* RBM-39 (Uniprot: Q9N368), *S. ratti* RBM-39 (Uniprot: A0A090LFF6), *C. briggsae* RBM-39 (Uniprot: A8XIX5), *C. japonica* RBM-39 (Uniprot: A0A4C1ZPV4), *C. remanei* RBM-39 (Uniprot: E3MXT8), *B. malayi* RBM-39 (Uniprot: A0A4E9ESP8), *T. muris* RBM-39 (Uniprot: A0A5S6R6A6), *X. tropicalis* RBM-39 (Uniprot: Q566M5), *R. norvegicus* RBM-39 (Uniprot: Q5BJP4), *P. troglodytes* RBM-39 (Uniprot:), *S. pombe* RBM-39 (Uniprot: O13845), *M. musculus* RBM-39 (Uniprot: Q8VH51), *H. sapiens* RBM39 (Uniprot: Q14498), *G. gallus* RBM-39 (Uniprot: E1BRU3), *D. melanogaster* Caper (Uniprot: Q9VM49), *B. taurus* RBM-39 (Uniprot: A0A3Q1LWZ4), *A. gambiae* RBM-39 (Uniprot: Q7PN29), *D. rerio* RBM-39 (Uniprot: Q58ER0), *C. lupus* RBM-39 (Uniprot: E2R4L0) and *P. pacificus* RBM-39 (Uniprot: H3FJ10) were used in Figure S2N.

### CRISPR/Cas9 mutant and reporter strains

In order to generate HA tagged *rnp-6* strains, guide RNAs were selected by using the web tool (https://zlab.bio/guide-design-resources). sgRNAs were synthesized with EnGen® sgRNA Synthesis Kit (NEB, #E3322) by following manufacturer’s protocol. CRISPR/Cas9 insertion was generated by following a co-CRISPR strategy ^49^. *dpy-10* was used as marker to enrich potential hits. Ribonucleoprotein complexes containing sgRNA, Cas9 and repair templates were annealed at 37°C for 15 minutes prior to injection. The primers used in this study are listed in Supplementary Table 13. For the GFP tagged RNP-6 strains, mKate2 tagged RBM-39 strains, mNeonGreen tagged EGL-8 strain and *rbm-39(S294L)* mutant strain, they were generated by Sunybiotech (https://www.sunybiotech.com). All the strains were validated by sanger sequence.

### Life span

All life spans were performed at 20°C unless otherwise noted. Worms were allowed to grow to the young adult stage on standard NGM plates with OP50. For each genotype, ~150 young adults were transferred to NGM plates with OP50 supplemented with 10 μM of FUdR. Survival was monitored every other day. Worms which did not respond to gentle touch by a worm pick were scored as dead and were removed from the plates. Animals that crawled off the plate or had ruptured vulva phenotypes were censored. All life span experiments were done at least three times independently unless otherwise noted. Graphpad Prism was used to plot survival curves. Survival curves were compared and p values were calculated using the log-rank (Mantel-Cox) analysis method. Complete life span data are available in Supplementary Table 16.

### Infection assay

S. aureus (Strain MW2) was grown in TSB medium at 37°C with gentle shaking overnight. 100 μl of the bacterial culture was seeded and spread all over the surface of the TSA plate with 10 μg/mL nalidixic acid (NAL). The plates were allowed to grow overnight at 37°C. On the next day, the plates were left at room temperature for at least 6 hours before the infection experiments. Around 25 synchronized young adult worms were transferred to the plates. Three technical replicate plates were set up for each condition. Worms were treated with 100 μM FUdR from L4 stage to prevent internal hatching during experiments. The plates were then incubated at 25°C to initiate infection experiment. Scoring was performed every day. Worms were scored as dead if the animals did not respond to gentle touch by a worm pick. Worms that crawled off the plates or had ruptured vulva phenotypes were censored from the analysis. All *C. elegans* killing assays were performed three times independently unless otherwise stated. Genotypes were blinded for all *C. elegans* infection survival experiments in order to eliminate any investigator-induced bias. Results of each biological replicate of infection survival experiments can be found in Supplementary Table 17.

### RNA interference in *C. elegans*

RNAi experiments were performed as previously described ^11^. *E. coli* HT115 and *E. coli OP50(xu363)* bacterial strains were used in this study. The HT115 bacteria were from the Vidal or Ahringer library. The *OP50(xu363)* competent bacteria were transformed with dsRNA expression plasmids which were extracted from the respective HT115 bacterial strains. The RNAi bacteria were grown in LB medium supplemented 100 μg/ mL ampicillin at 37 °C overnight with gentle shaking. The culture was spread on RNAi plates, which are NGM plates containing 100 μg/mL ampicillin and 0.4 mM isopropyl ß-D-1-thiogalactopyranoside (IPTG). RNAi expressing bacteria were allowed to grow on the plates at room temperature for two days. RNAi was initiated by letting the animals to feed on the desired RNAi bacteria. RNAi experiments related to Fig. 1d–f and Extended Data Fig. 2g-h were done with *OP50(xu363)* bacterial. For RNAi life span experiments related to Fig. 3a, m and n, worms were grown on HT115 RNAi bacterial from egg until day 1 adulthood and then transferred to NGM plates seeded with OP50.

### RNA extraction and cDNA synthesis

*C. elegans* were lysed with QIAzol Lysis Reagent. RNA was extracted using chloroform extraction. The samples were then purified using RNeasy Mini Kit (Qiagen). Purity and concentration of the RNA samples were assessed using a NanoDrop 2000c (peqLab). cDNA synthesis was performed using iScript cDNA synthesis kit (Bio-Rad). Standard manufacturers protocols were followed for all mentioned commercial kits.

### RNAseq and bioinformatic analysis

1 μg of total RNA was used per sample for library preparation. The protocol of Illumina Tru-Seq stranded RiboZero was used for RNA preparation. After purification and validation (2200 TapeStation; Agilent Technologies), libraries were pooled for quantification using the KAPA Library Quantification kit (Peqlab) and the 7900HT Sequence Detection System (Applied Biosystems). The libraries were then sequenced with Illumina HiSeq4000 sequencing system using paired end 2×100 bp sequencing protocol. For data analysis, Wormbase genome (WBcel235_89) was used for alignment of the reads. This was performed with the Hisat version 2.0.4 ^50^. Differentially expressed genes (DEGs) between different samples were identified using the Stringtie (version 1.3.0) ^51^, followed by Cufflinks (version 2.2) ^52^. The DAVID database ^53^ was used for enrichment and Gene Ontology (GO) analysis. SAJR pipeline ^54^ was used for splicing analysis. For intron retention analysis, Bedtools coverage (version 2.29.0) was used to count intron and total gene expression. IBB (version 20.06; R version 4.0.3) ^55^ was used to calculate differential intron expression. DCC/CircTest pipeline ^56^ was performed to quantify Circular RNAs expression. q value <0.05 is considered to be significant for SAJR and DEG analysis; p value <0.001 is considered to be significant for intron retention analysis.

### Alternative splicing PCR assay

Phusion Polymerase (Thermo Fisher) was used to amplify the *egl-8, tos-1 and tcer-1* segments. PCR reactions were cycled 30 times with an annealing temperature of 53 °C. The products were visualized by staining with Roti-GelStain (Carl Roth) after agarose gel electrophoresis. For primer sequences, please refer to Supplementary Table 15.

### Western blot

For *C. elegans* samples, animals were first washed with M9 buffer. Worm pellets were resuspended in RIPA buffer supplemented with cOmplete Protease Inhibitor (Roche) and PhosSTOP (Roche) and snap frozen in liquid nitrogen. Thawed samples were lysed using Bioruptor Sonication System (Diagenode). Protein samples were then heated to 95°C for 10 min in Laemmli buffer with 0.8% 2-mercaptoethanol in order to denature proteins. Samples were loaded on 4–15% Mini PROTEAN TGXTM Precast Protein Gels (Bio-Rad), and electrophoresis was performed at constant voltage of 200V for around 40 min. After separation, the proteins were transferred to PVDF membranes using Trans-Blot TurboTM Transfer System (BioRad). 5% bovine serum albumin (BSA) or 5% milk in Tris-buffered Saline and Tween20 (TBST) were used for blocking of the membranes. After antibody incubations and washing with TBST buffer, imaging of the membranes was performed with ChemiDoc Imager (BioRad). Western Lightning Plus Enhanced Chemiluminescence Substrate (PerkinElmer) was used as the chemiluminescence reagent. A list of antibodies is provided in Table 18.

For immunoblotting analyses using HEK293FT samples, cells were washed once in-well with serum-free DMEM, to remove FBS, and lysed in 250μl lysis buffer (50 mM Tris pH 7.5, 1% Triton X-100, 150 mM NaCl, 50 mM NaF, 2 mM Na-vanadate, 0.011 gr/mL β-glycerophosphate, 1x PhosSTOP phosphatase inhibitors, 1x cOmplete protease inhibitors) for 10 min on ice. Samples were clarified by centrifugation (14,000 x g, 15 min, 4 °C) and supernatants were transferred to new tubes. Protein concentration was measured using a Protein Assay Dye Reagent (Bio-Rad). Protein samples were subjected to electrophoretic separation on SDS-PAGE and analyzed by standard Western blotting techniques. In brief, proteins were transferred to nitrocellulose membranes (Amersham), stained with 0.2% Ponceau solution (Serva) to confirm equal loading. Membranes were blocked with 5% powdered milk in PBS-T (1x PBS, 0.1% Tween-20) for 1 hour at room temperature, washed 3x 10 min with PBS-T and incubated with primary antibodies (1:1,000 in PBS-T, 5% BSA) rotating overnight at 4°C. The next day, membranes were washed 3x 10 min with PBS-T and incubated with appropriate HRP-conjugated secondary antibodies (1:10,000 in PBS-T, 5% milk) for 1 hour at RT. Signals were detected by enhanced chemiluminescence (ECL), using the ECL Western Blotting Substrate (Promega); or SuperSignal West Femto Substrate (Thermo Fisher Scientific) for weaker signals. Immunoblot images were captured on films (GE Healthcare).

### Co-immunoprecipitation

Worms expressing HA∷RNP-6, RBM-39∷mKate2 or both were harvested, and proteins were extracted by following standard protocol ^57^. A solubilization buffer containing 0.5% NP40, 150 mM NaCL and 50 mM Tris pH 7.4 supplemented with cOmplete Protease Inhibitor (Roche) and PhosSTOP (Roche) was used for immuno-precipitation. Flag immunoprecipitation was performed using Dynabeads Protein G (ThermoFisher Scientific) and FLAG M2 mouse monoclonal antibody (Sigma), following manufacturer’s protocols. Proteins were eluted from the beads by boiling with Laemmli buffer.

### Worm imaging

Analysis of worm reporters GFP∷RNP-6, RBM-39∷mKate2, mNeonGreen∷EGL-8 and HLH-30∷mNeonGreen were performed on a Zeiss Axioplan2 microscope with a Zeiss AxioCam 506 CCD camera. Fiji software (2.0.0) ^58^ was used for quantifying fluorescent intensity. For mNeonGreen∷EGL-8 images, the head neuron region was selected for quantification. For HLH-30∷mNeonGreen images, the nucleus of hypodermal cells were selected for quantification. For GFP∷RNP-6 images, the whole worm was selected for quantification. To reduce bias, individual worms were randomly picked under a dissection microscope and imaged. At least 20 worms per genotype were picked for imaging and all the experiments were done three times.

### Transient knockdowns in HEK293FT cells (siRNA transfections)

Transient knockdowns were performed using a pool of 4 siGENOME siRNAs (Horizon Discoveries) against PUF60, while an RLuc duplex siRNA that targets the *R. reniformis* Luciferase gene (Horizon Discoveries) was used as control. In brief, HEK293FT cells were seeded in 12-well plates at 20% confluence and the following day transfected with 20 nM of the siRNA pool using Lipofectamine RNAiMAX (Thermo Fisher Scientific) according to manufacturer’s instructions. Cells were harvested or fixed 72 hours post-transfection and knock-down efficiency was verified by immunoblotting.

### Immunofluorescence and confocal microscopy in HEK293FT cells

Immunofluorescence / confocal microscopy experiments and quantification of colocalization were performed as previously described ^59^. In brief, cells were seeded on fibronectin-coated coverslips and treated as indicated in each experiment. After treatments, cells were fixed for 10 min at room temperature with 4% PFA in PBS. Samples were washed/permeabilized with PBT solution (1x PBS, 0.1% Tween-20), and blocked with BBT solution (1x PBS, 0.1% Tween-20, 0.1% BSA). Staining was performed with the indicated primary antibodies in BBT (1:200 dilution) for 2 hours at RT for mTOR and LAMP2 staining or overnight at 4°C for TFE3 staining. Next, samples were washed 4x with BBT (15 min each), followed by incubation with appropriate highly cross-adsorbed secondary fluorescent antibodies for 1 hour at RT. Finally, nuclei were stained with DAPI and cells mounted on slides using Fluoromount-G (Invitrogen). Images from single channel captures are shown in grayscale. For the merged images, Alexa 488 is shown in green, TRITC in red and DAPI in blue. Images were captured using a 40x objective lens on an SP8 Leica confocal microscope. To quantify colocalization of mTOR signal with the lysosomal marker LAMP2, the Fiji software (Version 2.1.0/1.53c) ^58^ was used to define regions of interest (ROIs) corresponding to individual cells, excluding the nucleus. Fifty (50) individual cells from five independent fields were selected for the analysis. The Coloc2 plugin was used to calculate the Manders’ colocalization coefficient (MCC), using automatic Costes thresholding ^60,61^. MCC yields the fraction of the mTOR signal that overlaps with the LAMP2 signal. Subcellular localization of TFE3 was analyzed by scoring cells based on the signal distribution of TFE3, as shown in the example images in Fig. 7C. Signal was scored as nuclear (more TFE3 signal in the nucleus) or cytoplasmic (similar TFE3 signal between nucleus and cytoplasm). Cells from 5 independent fields, containing approximately 70 individual cells, were scored per genotype for each experiment.

### Statistical analysis

In all figure legends, ‘n’ denotes the number of independent replicate experiments performed, while ‘N’ indicates the total number of animals analyzed in each condition. All statistical analyses were performed in GraphPad Prism. Asterisks denote corresponding statistical significance *p < 0.05; **p < 0.01; ***p < 0.001.

## Data availability

All RNA-seq datasets generated and analyzed in this study are available in the GEO datasets with the accession number PRJNA757629. All other data are available from the corresponding author upon request.

## Acknowledgements

We wish to thank the Caenorhabditis Genetics Center (University of Minnesota) and the Japanese National Biosource Project for providing *C. elegans* strains; Stephen Nurrish (Harvard Medical School) for sharing *egl-8* plasmids; Franziska Metge and Jorge Boucas from the MPI-AGE Bioinformatics, Christian Kukat from MPI-AGE Imaging and Xinping Li from MPI-AGE Proteomics Cores for technical support. We would also like to thank Orsolya Symmons, Balaji Srinivasan, Kazuto Kawamura and Shamsh Tabrez Syed for valuable comments on the manuscript. C.D. is funded by the European Research Council (ERC) under the European Union’s Horizon 2020 research and innovation programme (grant agreement No 757729). This work was supported by the Max Planck Society, Germany.

## Author Contributions

W.H. and A.A. conceived, designed the study. W.H. and A.L. performed the EMS mutagenesis, mapping experiments and life span experiments. W.H. and A.L. performed western blot experiments. W.H., C.K. and L.H. performed infection assay. C.K. performed Co-IP experiments. W.H., C.K. and A.L. prepared RNA samples for RNAseq. S.F. and C.D. designed and performed human cell culture experiments. W.H. and A.A. wrote the manuscript with input from all authors.

